# CTCF-binding element regulates ESC differentiation via orchestrating long-range chromatin interaction between enhancers and HoxA

**DOI:** 10.1101/2020.10.14.340174

**Authors:** Guangsong Su, Wenbin Wang, Jun Chen, Man Liu, Jian Zheng, Dianhao Guo, Jinfang Bi, Zhongfang Zhao, Jiandang Shi, Lei Zhang, Wange Lu

## Abstract

Proper expressions of Homeobox A cluster genes (*HoxA*) are essential for embryonic stem cells (ESC) differentiation and individual development. However, the mechanisms underlying controlling precise spatiotemporal expressions of *HoxA* during early ESC differentiation remain poorly understood. Herein, we identified a functional CTCF-binding element (CBE^+47^) closest to the 3’-end of *HoxA* within the same topologically associated domain (TAD) in ESC. CRISPR-Cas9 mediated the CBE^+47^ deletion significantly promotes retinoic acid (RA)-induced *HoxA* expression and early ESC abnormal differentiation. In delineating its underlying mechanisms, we find that CBE^+47^ can precisely organize chromatin interactions between its adjacent enhancers and *HoxA* by using chromosome conformation capture assay (Capture-C). Furthermore, we also find that its adjacent enhancers as enhancer-enhancer interaction complex (EEIC) are required for RA-induced *HoxA* activation. Collectively, these results provide a new insight for RA-induced *HoxA* expression, also highlight the unique and precise regulatory roles of CBE in ESC differentiation.

## INTRODUCTION

Embryonic stem cells (ESC) are cells derived from the inner cell mass (ICM) (*Martin, 1981*). ESC have the ability of unlimited self-renewal and differentiation into three embryonic germ layers (*Takahashi et al., 2007; Takahashi & Yamanaka, 2006*). Therefore, ESC have great application potential in regenerative medicine and tissue engineering (*Metcalfe & Ferguson, 2007; N. Sato, Meijer, Skaltsounis, Greengard, & Brivanlou, 2004*). Different state of ESC requires proper expression of specific gene. The pluripotency state of ESC requires expressions of master control transcription factors such as *Oct4/Nanog/Sox2* (*Boyer et al., 2005; Loh et al., 2006*), and expressions of genes such as *HoxA/Gata4/Pax6* are essential for ESC differentiation (*Fujikura et al., 2002; Gajovic, St-Onge, Yokota, & Gruss, 1997; Martinez-Ceballos, Chambon, & Gudas, 2005*). In recent years, studies have shown that high-order chromatin structures play a critical role in regulating gene expression and maintaining ESC status (*Levasseur, Wang, Dorschner, Stamatoyannopoulos, & Orkin, 2008; Wei et al., 2013*). Numerous studies have revealed that topologically associated domains (TAD) are the basic unit of nuclear chromatin (*Luppino et al., 2020; Pope et al., 2014*). Functional DNA elements, such as enhancers, are more likely to promote expression of target gene through directly long-range chromatin interactions within the same TAD (*Ji et al., 2016; Lupianez et al., 2015*). Long-range chromatin interactions depend on the proteins that mediate its interaction, among them CTCF plays the central role. Nora Elphege P et al. used auxin-induced degradation of CTCF protein and found that the absence of CTCF led to disappearance of TAD, revealing that CTCT is essential for maintaining the formation of TAD (*Nora et al., 2017*). CTCF forms chromatin-loop through directly binding the CTCF-binding elements (CBE) enriched at TAD boundary and relies on its binding polarity criteria to maintain the formation of TAD (*de Wit et al., 2015; Guo et al., 2015; Nanni, Ceri, & Logie, 2020*). Although many studies have found that the CBE at TAD boundary plays key regulatory roles in biological development, the research on CBE inside the TAD is relatively lacking.

Retinoic acid (RA) can induce ESC differentiation rapidly (*Bain, Kitchens, Yao, Huettner, & Gottlieb, 1995; Tay, Zhang, Thomson, Lim, & Rigoutsos, 2008*). In this process, *HoxA* are significantly activated (*De Kumar et al., 2015; Kashyap et al., 2011*). Recent studies have shown that CBE plays the core regulatory roles in tumorigenesis (*Li et al., 2020; Huacheng Luo et al., 2018*), immunogenesis (*Ba et al., 2020; Jain, Ba, Zhang, Dai, & Alt, 2018*) and stem cell development (*Arzate-Mejia, Recillas-Targa, & Corces, 2018; H. Zheng & Xie, 2019*) by controlling target genes expression through organizing high-order chromatin structures. We and other research groups have found that multiple enhancers are required for RA-induced *HoxA* expression and early ESC differentiation by long-range chromatin interactions with *HoxA* (*Cao et al., 2017; Liu et al., 2016; Su et al., 2019; Yin et al., 2015*). Besides, previous studies also have reported many regulatory patterns for *HoxA* activation during ESC differentiation, among them, several CBE (also known as CBS, CTCF-binding site) within *HoxA* locus can organize *HoxA* chromatin structures, and play significant regulatory roles in ESC differentiation and leukemogenesis (*Ghasemi, Struthers, Wilson, & Spencer, 2020; Huacheng Luo et al., 2018; Narendra et al., 2015*). However, it is unknown whether CBE plays functional roles in regulating *HoxA* expression by orchestrating enhancer chromatin structures during RA-induced early ESC differentiation.

In this study, we identified a new functional CTCF-binding element (CBE^+47^) required for RA to activate *HoxA* expression and promote early ESC proper differentiation. On the mechanisms, our results show that CBE^+47^ can regulate the interactions between multiple enhancers, as well as the interactions of these enhancers with *HoxA* chromatin. Our study reveals that a new functional CBE^+47^ can regulate the precise expressions of *HoxA* by accurately organizing chromatin interactions between its adjacent enhancers and *HoxA*, thereby facilitating early correct differentiation of RA-induced ESC. These findings not only increase our understanding of mechanisms of RA-induced *HoxA* activation and early ESC differentiation, but also highlight the unique and precise regulatory roles of CBE in high-order chromatin structures.

## RESULTS

### Identification of a CTCF-binding element (CBE^+47^) closest to the 3’-end of *HoxA* within the same TAD in ESC

The basic structural unit in high-order chromatin structure is the TAD, and DNA functional elements such as enhancers or insulators in the same TAD are more likely to regulate the precise expression of neighboring genes (*Ji et al., 2016; Lupianez et al., 2015*). To discover potential DNA functional elements that regulate the expression of *HoxA*, we first analyzed the Hi-C data in ESC. We find that 5’-end of *HoxA* is located at the TAD boundary region, and 3’-end is within the TAD (*Figure 1A*). Previous studies have shown that several significant CTCF-binding elements (CBE) are located at the *HoxA* locus, and play an important regulatory role in *HoxA* expression and ESC differentiation (*Narendra et al., 2015; Rousseau et al., 2014*). Interestingly, many significant CBE are also found in the same TAD at 3’-end of *HoxA* (*Figure 1B*), but there are no reports on whether these CBE have regulatory effects on *HoxA* expression and ESC differentiation.

**Figure 1.**
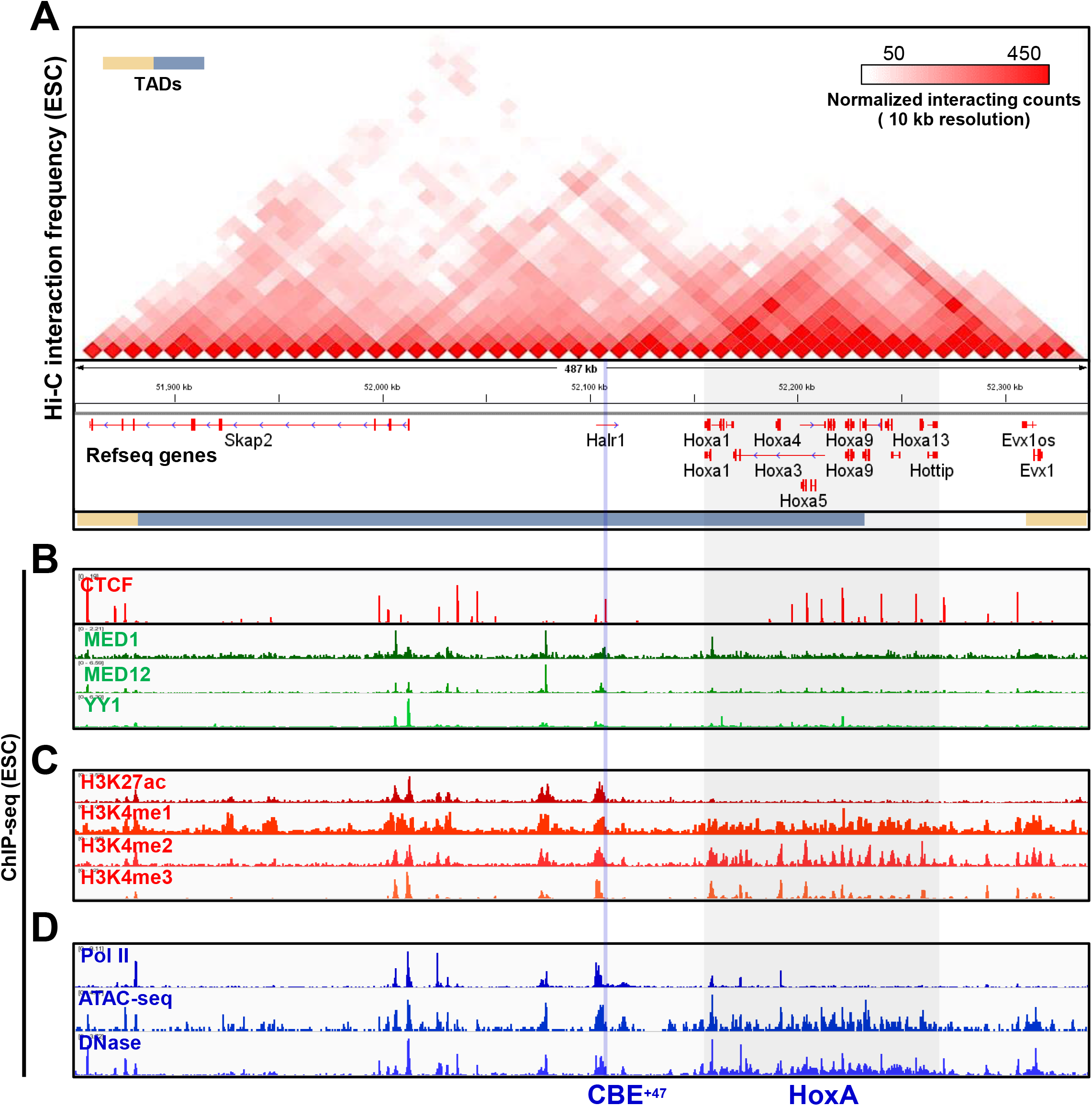
Identification of a CBE (CBE^+47^) closest to the 3’-end of *HoxA* within the same TAD in ESC. (**A**) Hi-C interaction map of the ~0.5 Mb region surrounding the *HoxA* in undifferentiated ESC. Data were extracted from Bonev B et al. 2017. (**B**) IGV (Integrative Genomics Viewer) screenshots showing gene tracks of CTCF, MED1, MED12 and YY1 ChIP-seq signal occupancy at Skap2 and *HoxA* loci in ESC. CTCF ChIP-seq shows CTCF binding throughout the locus. Multiple CTCF sites are located at the downstream of *HoxA* locus. (**C, D**) IGV view of selected ChIP-seq tracks at Skap2 and *HoxA* loci in ESC. Shown are H3K27ac, H3K4me1, H3K4me2, H3K4me3, Pol ll, ATAC-seq and DNase. Blue shadowing indicates the CBE^+47^ region. Grey shadowing shows the *HoxA* locus.

Here, we focus on a significant CBE closest to the 3’-end of *HoxA*, about 47 kb from *HoxA* (CBE^+47^), located in the second intron of *Halr1* (*Figure 1B; Figure 1–figure supplement 1*). We find significant binding of Med1, Med12 and YY1 at upstream of CBE^+47^ (*Figure 1B*). The epigenetic modification-related markers (e.g. H3K27ac, H3K4me1/2/3) are also enriched upstream of CBE^+47^ (*Figure 1C*). Furthermore, we also find significant chromatin accessibility (DNase and ATAC-seq) and RNA transcription activity (PolII) enrichment in these same regions (*Figure 1D*). These data indicate that CBE^+47^ is located between the potential regulatory elements and *HoxA*. Previous studies have shown that CBE can precisely regulate target gene expression through organizing chromatin interactions between functional element and target gene (*Hark et al., 2000; Kurukuti et al., 2006*). Therefore, based on the above results, we speculate that CBE^+47^ may have a regulatory role.

### Genes expression analysis in WT and CBE^+47^ deletion cells under self-renewal culture condition

To investigate whether CBE^+47^ has a regulatory effect, CRISPR-Cas9 mediated deletion method was used for further research (*Su et al., 2019*). Two sgRNAs were designed in upstream and downstream of CBE^+47^ (*Figure 2A*), after Cas9 cleavage and DNA recombination, we obtained two homozygous CBE^+47^ knockout cell lines (CBE^+47^-KO). Genomic DNA PCR using specific primer, and Sanger sequencing confirmed the deletion of CBE^+47^ (*Figure 2B*). AP-staining showed that there were no significant differences in cell morphology in the CBE^+47^ knockout cell lines under self-renewing culture conditions (*Figure 2C*). qRT-PCR results showed that mRNA expression levels of *Halr1* and *Sakp2* did not change significantly (*Figure 2D*). However, 3’-end genes of *HoxA* were significantly up-regulated (*Hoxa2-a6*), while TAD boundary gene *Hoxa9* was significantly down-regulated in CBE^+47^-KO cells compared with WT (*Figure 2E*). These data suggest that deletion of CBE^+47^ significantly changes expressions of *HoxA*, indicating that even in undifferentiated ESC state, the CBE^+47^ is also required to maintain proper expressions of *HoxA*.

**Figure 2.**
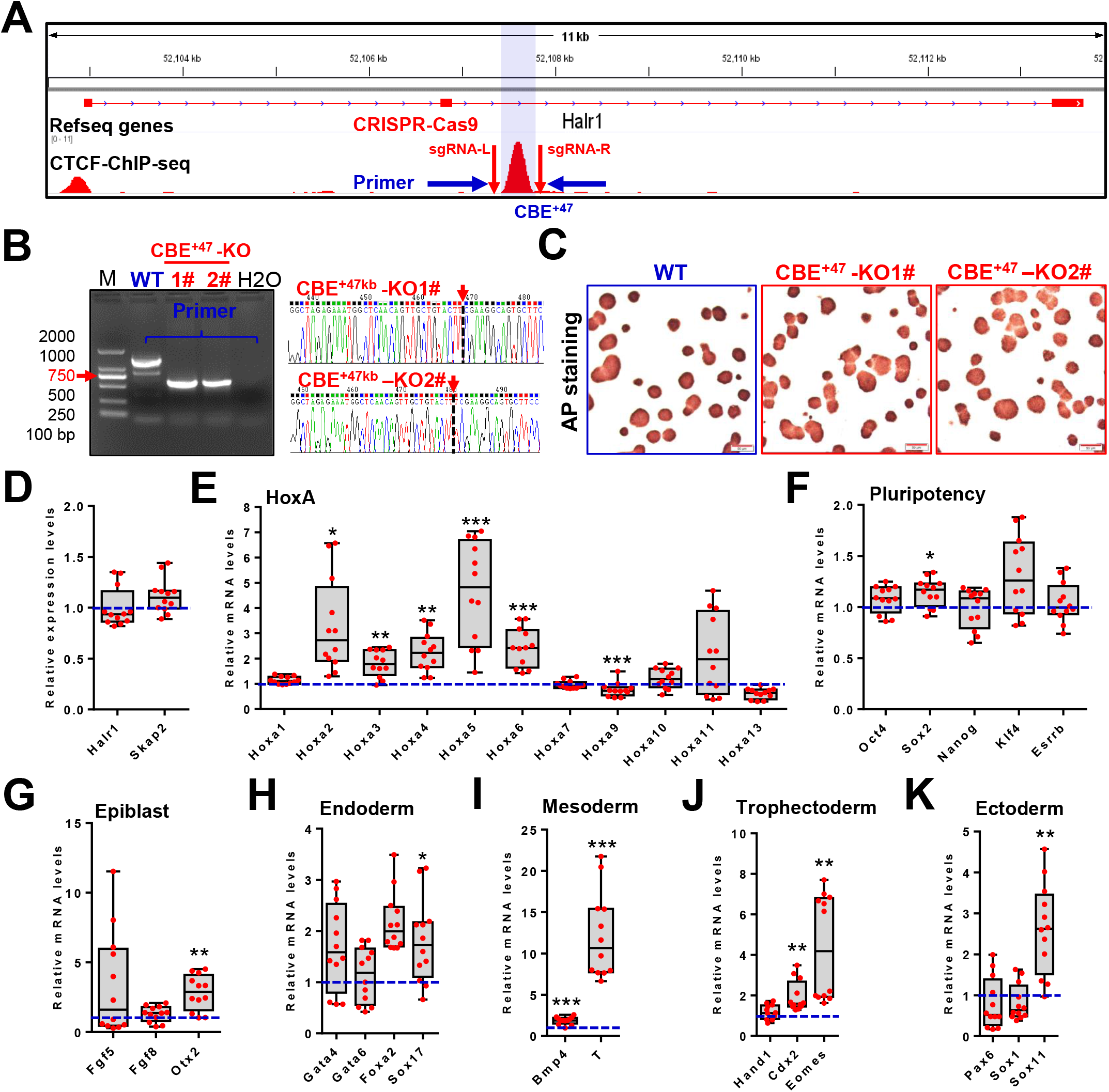
Gene expression analysis in WT and CBE^+47^ deletion cells under self-renewal culture condition. (**A**) Schematic showing CRISPR/Cas9-mediated deletion of the CBE^+47^ (blue shadowing) using two sgRNAs. Indicated primer was used to distinguish CBE^+47^-KO (CBE^+47^ knockout) from wild-type (WT) clones. (**B**) Validation of knockout cell lines by genomic DNA PCR and sequencing. Shown are images from representative clones. DNA sequencing of two CBE^+47^-KO cell clones (#1 and #2) using the primer. (**C**) Overview appearance of WT and CBE^+47^-KO cells with alkaline phosphatase (AP) staining. (**D-K**) qRT-PCR of WT and CBE^+47^-KO cells showing mRNA expression of *Skap2* and *Halr1* (**D**), *HoxA* (**E**), pluripotency genes (**F**), epiblast genes (**G**), endoderm genes (**H**), mesoderm genes (**I**), trophectoderm genes (**J**) and ectoderm genes (**K**) in undifferentiated ESC. In (**D-K**), data are represented as min to max. The expression level in WT ESC was relatively set to 1 (As shown in blue dotted line). Indicated significance is based on Student’s *t*-test (Two-tailed, * indicates *P*<0.05, ** indicates *P* <0.01, *** indicates *P* < 0.001). In (**D-K**), n = 12, including 2 CBE^+47^-KO cell lines (CBE^+47^-KO1# and CBE^+47^-KO2#), and 2 biological replicates per cell line with 3 technical replicates in 1 biological replicate. In (**B**), M: DNA Marker. In (**C**), scale bar, 50 μm. Source files are available in *Figure 2—source data 1*.

Considering that expression of *HoxA* is critical for ESC differentiation (*De Kumar et al., 2017; Martinez-Ceballos et al., 2005; Yin et al., 2015*), and CBE^+47^ deletion significantly increased some *HoxA* expression, we then tested if the ESC’s pluripotency status was impacted by examining the expressions of pluripotency- and differentiation-related genes by qRT-PCR (*Figure 2F-K*). No significant differences in the expressions of pluripotency-regulated genes, except for a slight up-regulation of *Sox2*. Mesoderm and trophectoderm genes are significantly up-regulated. Expression of other germ layer genes such as *Otx2* (epiblast), *Sox17* (endoderm) and *Sox11* (ectoderm) also significantly increased. These results indicate that deletion of CBE^+47^ does not regulate the pluripotency of ESC, but may affect its differentiation ability.

### CBE^+47^ deletion significantly promotes RA-induced *HoxA* expression and early ESC abnormal differentiation

During the early differentiation of ESC induced by RA, *HoxA* are significantly activated compared with undifferentiated ESC (*De Kumar et al., 2015; Yin et al., 2015*) (*Figure 3–figure supplement 1*). Therefore, we would like to know whether the CBE^+47^ plays a regulatory role in regulating *HoxA* in the early stage of RA-induced ESC differentiation. ESC differentiation was induced by RA to 3 different time points (*Figure 3–figure supplement 2A*). qRT-PCR was used to analyze expression levels of *HoxA*, and the results showed that *HoxA* 3’-end genes had significantly higher expressions (*Hoxa2-a6*), and TAD boundary genes (*Hoxa7* and *Hoxa9*) also show significantly increased expression levels. However, *HoxA* 5’-end genes did not change significantly (*Hoxa10, Hoxa11* and *Hoxa13*) (*Figure 3A-C*). In addition, we also find that antisense long non-coding RNAs at *HoxA* locus are also significantly up-regulated, such as *Hoxaas3* (*Figure 3–figure supplement 2B-D*). These data indicate that CBE^+47^ knockout significantly promotes RA-induced expression of *HoxA*, revealing that the CBE^+47^ is to restrict RA-induced over-expression of *HoxA*.

**Figure 3.**
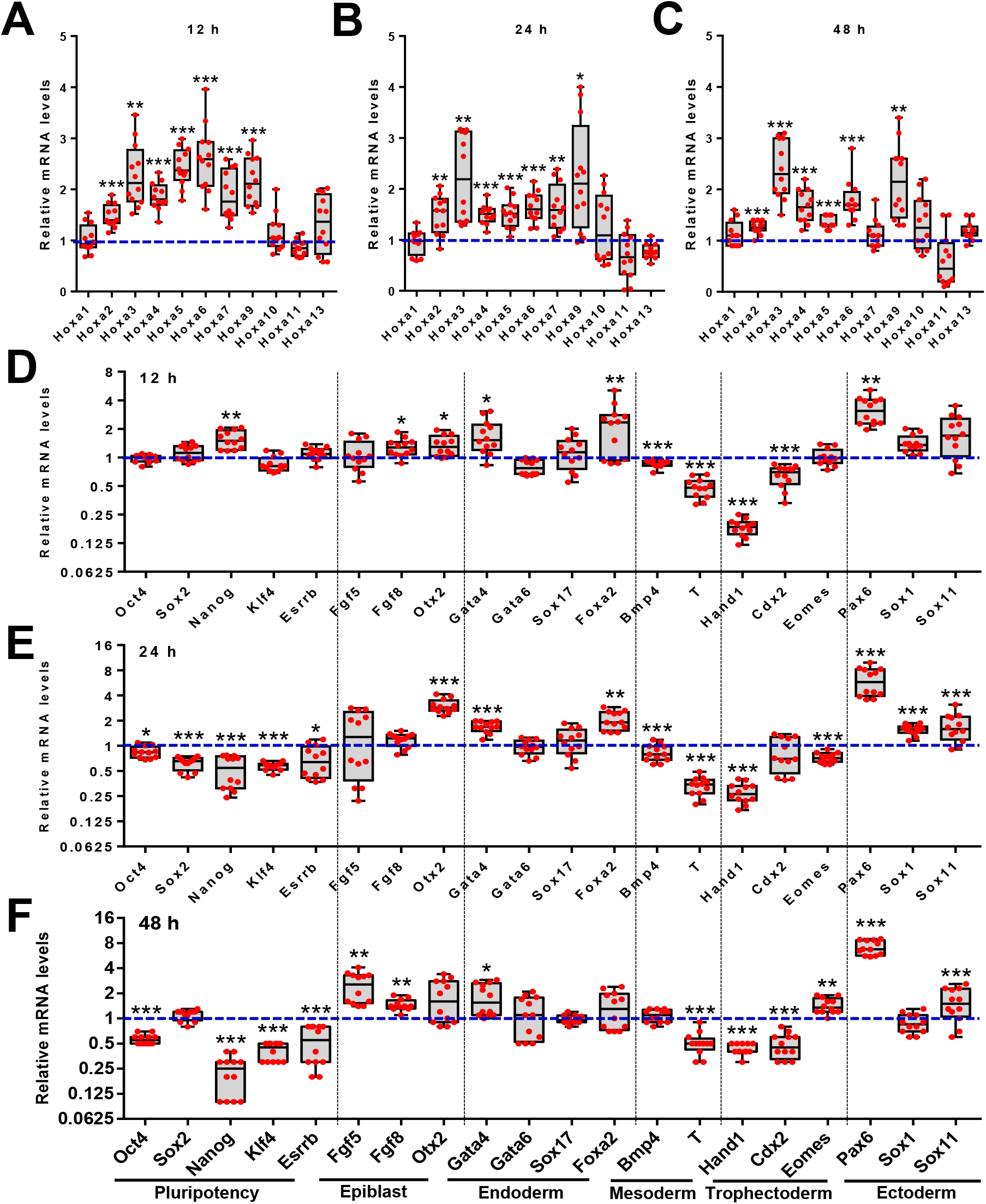
CBE^+47^ deletion significantly promotes RA-induced expressions of *HoxA* and early ESC abnormal differentiation. (**A-C**) qRT-PCR of WT and CBE^+47^-KO cells showing expressions of *HoxA* in ESC after RA treatment (12h, 24h and 48h). (**D-F**) qRT-PCR of WT and CBE^+47^-KO ESC showing mRNA expressions of pluripotency genes, epiblast genes, endoderm genes, mesoderm genes, trophectoderm genes and ectoderm genes in RA-induced ESC (12h, 24h and 48h). In (**A-F**), data are represented as min to max. The expression level in WT ESC was relatively set to 1 (As shown in blue dotted line). Indicated significance is based on Student’s *t*-test (Two-tailed, * indicates *P* < 0.05, ** indicates *P* < 0.01, *** indicates *P* < 0.001). In (**A-F**), n = 12, including 2 CBE^+47^-KO cell lines (CBE^+47^-KO1# and CBE^+47^-KO2#), and 2 biological replicates per cell line with 3 technical replicates in 1 biological replicate. Source files are available in *Figure 3—source data 1*.

Furthermore, to investigate whether CBE^+47^ knockout affects RA-induced ESC differentiation, we used the qRT-PCR method to analyze the changes of pluripotency- and differentiation-related master control genes in WT and CBE^+47^-KO ESC under RA condition. The results show that the expressions of the pluripotency master control genes did not change significantly at 12 hours after ESC differentiation but significantly decreased at 24 and 48 hours (such as *Nanog* and *Klf4*). Epiblast master control gene (*Otx2*), endoderm master control gene (*Gata4*) and ectoderm master control gene (*Pax6*) have significantly high expression levels, while mesoderm master gene (*T*) and trophectoderm master gene (*Hand1*) are significantly down-regulated, these data suggesting that deletion of CBE^+47^ significantly leads to ESC abnormal differentiation (*Figure 3D-E*). Combining these results reveals that the CBE^+47^ is essential to maintain RA-induced *HoxA* expression and early ESC proper differentiation.

### Transcriptome analysis in WT and CBE^+47kb^ deletion cells during RA-induced early ESC differentiation

To further investigate the impacts of CBE^+47^ knockout on RA-induced early ESC differentiation (24h), we performed transcriptome analysis by RNA sequencing (RNA-seq) after RA-induced differentiation of WT and CBE^+47^ knockout cells. Compared with WT cells, a total of 869 genes were up-regulated and 360 genes down-regulated in CBE^+47^-KO cells after statistical analysis (Fold change ≥ 2, *P* < 0.05) (*Figure 4A*). Among them, *HoxA* are significantly up-regulated (*Figure 4B*). Because the *HoxA* 3’-end genes located in the same TAD with CBE^+47^, compared with those genes in two adjacent TADs, we find that CBE^+47^ deletion significantly promotes gene expression in the same TAD, while genes in two adjacent TADs have no significant differences, indicating that direct regulation of CBE^+47^ is restricted within the TAD. These findings reveal that CBE^+47^ services as a *cis*-regulatory element playing a local regulatory role (*Figure 4–figure supplement 1*). Moreover, compared with WT cells, we also find that *HoxB/C/D* cluster genes also have significantly higher expressions (*Figure 4–figure supplement 2A*). Relevant genes in the RA signaling pathway are also significantly over-expressed, such as *Crabp2, Cyp26a1* and *Stra6*.

**Figure 4.**
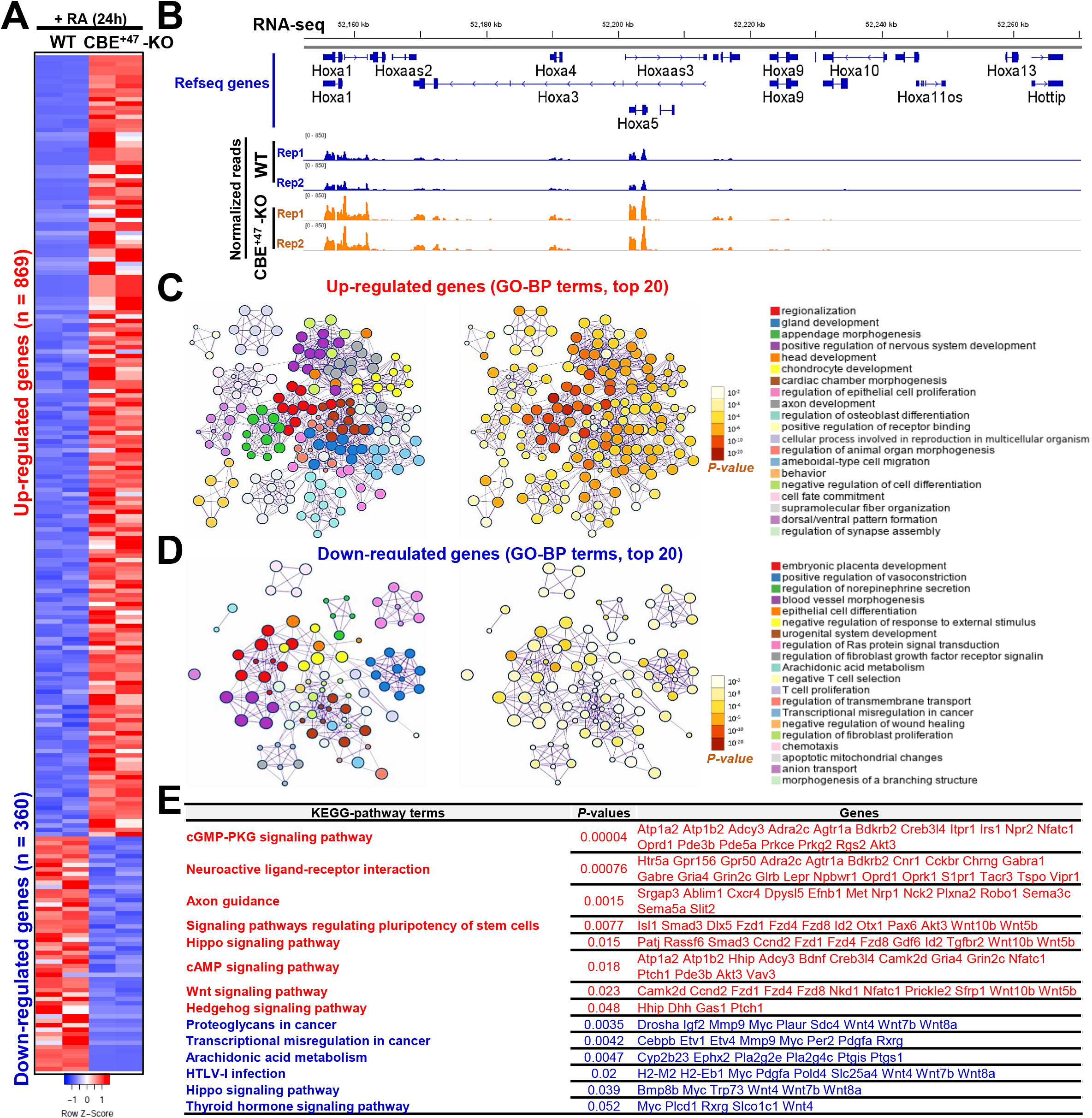
Transcriptome analysis in WT and CBE^+47^ deletion cells during RA-induced early ESC differentiation. (**A**) Heatmap depicting gene expression changes in WT and CBE^+47^-KO cells treated with RA for 24 h (Fold-change ⩾ 2, and *P* < 0.05, as determined by DESeq2). (**B**) RNA-seq results shown the Heatmap of *HoxA*. (**C-D**) GO-BP and KEGG pathway analyses indicated differentially expressed genes. In (**E**), the red font shows the signaling pathways of up-regulated genes enrichment, and the blue font shows the signaling pathways of down-regulated genes enrichment. Source files are available in *Figure 4—source data 1*.

Differentiation-related genes *Cbx4* and *Foxa1* are also significantly up-regulated (*Figure 4–figure supplement 2B*). On the contrary, we also find that genes related to pluripotency such as *Myc, Lin28a* and *Etv4* are significantly suppressed (*Figure 4-figure supplement 2C*). These data indicate that in early differentiation of RA-induced ESC, CBE^+47^ deletion significantly promotes early ESC differentiation, suggesting that the CBE^+47^ may inhibit over-differentiation of ESC, thereby ensuring its accurate normal differentiation.

In addition, to better understand the molecular mechanisms of CBE^+47^ regulatory roles during RA-induced early ESC differentiation, we performed bioinformatic analysis for the up- and down-regulated genes (*Figure 4–figure supplement 3*). GO analysis indicated that biological processes (BP) of up-regulated genes are mainly involved in gland development, positive regulation of nervous system development, etc (*Figure 4C*). Also, GO-BPs of down-regulated genes are mainly involved in embryonic placenta development, positive regulation of vasoconstriction, epithelial cell differentiation, etc (*Figure 4D*). KEGG pathway enrichment analysis reveals that up-regulated genes are mainly enriched in pathways including *cGMP-PKG* signaling pathway, neuroactive ligand-receptor interaction, axon guidance, signaling pathways regulating pluripotency of stem cells, *Hippo* signaling pathway, *cAMP* signaling pathway, *Wnt* signaling pathway and Hedgehog signaling pathway (*Figure 4E*). Enrichment results of down-regulated genes show that most pathways are related to proteoglycans in cancer, Transcriptional misregulation in cancer, Arachidonic acid metabolism, HTLV-I infection, *Hippo* signaling pathway and Thyroid hormone signaling pathway (*Figure 4E*). These results suggest that CBE^+47^ deletion leads to abnormal genes expression, indicating the essential role of CBE^+47^ for RA-induced early ESC proper differentiation.

### CBE^+47^ is required for proper chromatin interactions between its adjacent enhancers and *HoxA*

Previous studies have reported that CTCF regulates target gene expression through controlling chromatin interaction between enhancer and target gene, and this pattern occurs within a TAD (*Zhao et al., 2020*). Studies by us and others have found that there are three enhancers at the 3’-end of *HoxA* (*Figure 5A*). The three enhancers (e.g., E1, E2 and E3) are required for *HoxA* expression during RA-induced ESC differentiation (*Cao et al., 2017; Liu et al., 2016; Su et al., 2019; Yin et al., 2015*). Among them, it has been reported that E1 and E3 have significant long-range interactions with *HoxA* chromatin. Remarkably, CBE^+47^ is just located between the three enhancers and *HoxA* (*Figure 5A*). Therefore, we speculate that this CBE may be involved in organizing the interactions between the three enhancers and *HoxA* chromatin.

**Figure 5.**
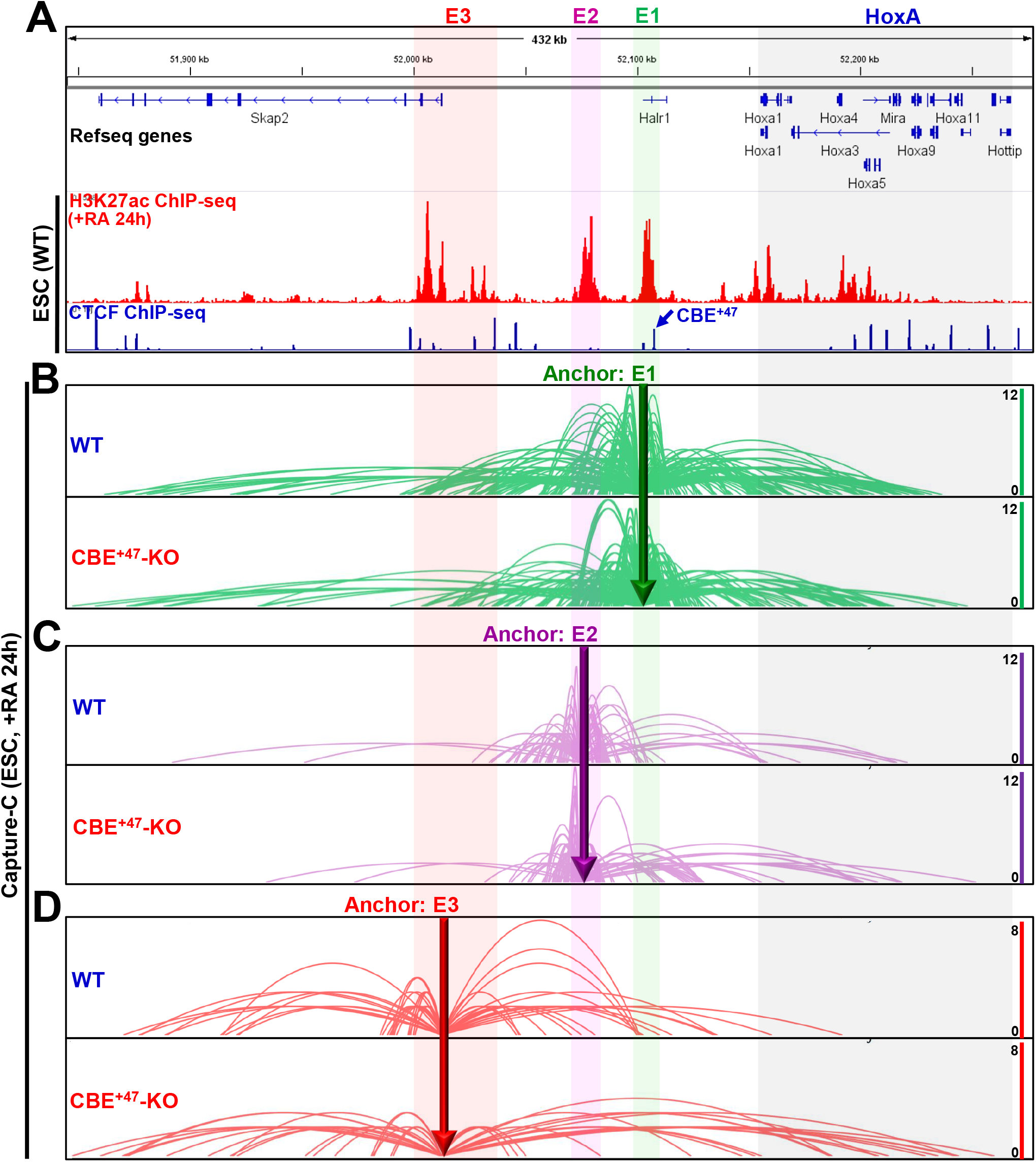
The CBE^+47^ is required for proper chromatin interactions between its adjacent enhancers and *HoxA*. (**A**) IGV (Integrative Genomics Viewer) screenshots showing gene tracks of H3K27ac (+RA, 24h) and CTCF ChIP-seq signal occupancy at *Skap2* and *HoxA* loci in WT ESC. (**B-D**) Chromatin interaction profiles by using enhancers as the anchors in WT and CBE^+47^-KO cells under RA treatment for 24h. Y-axis denotes interaction frequency represented by the number. Light green, light purple, light red and light grey shadows mark the E1, E2, E3 and *HoxA* regions, respectively.

Many experimental techniques, such as Capture-C, have been widely used to study the precise chromatin interactions between enhancers and target genes (*Davies et al., 2016; Hughes et al., 2014; Liang et al., 2020; Qian et al., 2019; Su et al., 2019; Zhang et al., 2020*). We used the Capture-C method to study different interactions between enhancers and *HoxA* chromatin in WT and CBE^+47^-KO cells. We used cells induced by RA for 24h to carry out Capture-C, E1, E2 and E3 as the anchor regions, respectively (*Figure 5B-D*). The results show that in WT cells, there are significant interactions between enhancers, and these enhancers also interact with *HoxA* chromatin. Among them, E1 interacts with *HoxA* more strongly (*Figure 5B*). Compared with WT cells, the interactions between enhancers are significantly reduced in CBE^+47^ knockout cells. However, E2 and E3 interact with *HoxA* chromatin increased significantly, and the intensity of chromatin interactions shifted from 3’-end to 5’-end of *HoxA* (*Figure 5C, D*). These results indicate that the deletion of CBE^+47^ significantly affects interactions between enhancers and *HoxA* chromatin, revealing that the CBE^+47^ is essential to maintain these proper chromatin interactions during RA-induced early ESC differentiation.

### Multiple enhancers deletion shows synergistic effect on RA-induced *HoxA* expression

To gain a deeper understanding of the regulatory effects of these enhancers on *HoxA* in our system, once again, we used the CRISPR-Cas9 method to knock out these enhancers. We obtained multiple homozygous enhancer knockout cell lines (ie ΔE1, ΔE2, ΔE3, ΔE1/2, ΔE1/2/3) (*Figure 6–figure supplement 1*). After RA treatment for 24 hours, qRT-PCR was used to determine expressions of *HoxA* (*Figure 6A-E*). The results show that in ΔE1 and ΔE2 cells, enhancers knockout have a similar effect on the regulation of *HoxA*, that is, both the 3’-end (*Hoxa1*) and the central genes (*Hoxa5-a10*) are significantly suppressed (*Figure 6A, B*); In ΔE3 cells, *Hoxa1-a7* are significantly inhibited, which is significantly different from ΔE1 and ΔE2, indicating that these enhancers have their specific regulatory effects (*Figure 6C*). While, in ΔE1/2 and ΔE1/2/3 cells, expressions of *HoxA* are significantly reduced compared with ΔE1 or ΔE2 or ΔE3 (*Figure 6D-E; Figure 6–figure supplement 2*). The above results demonstrate that these enhancers are required for RA-induced *HoxA* expression, indicating that the interaction among these three enhancers and *HoxA* chromatin showing a synthetic regulatory effect, which in turn promotes RA-induced *HoxA* activation.

**Figure 6.**
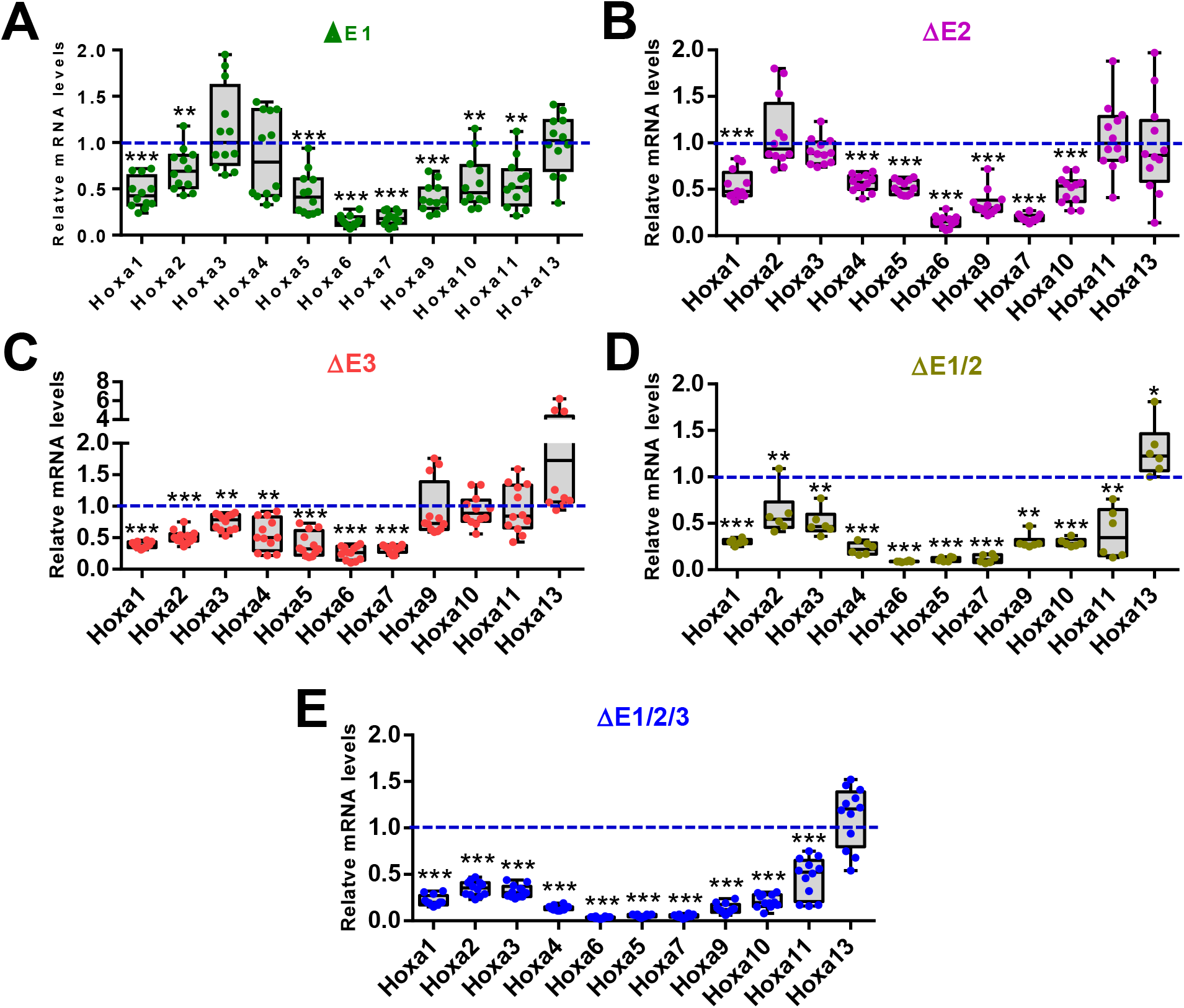
Multiple enhancers deletion shows synergistic effect on RA-induced *HoxA* expression. *HoxA* mRNA levels were measured by qRT-PCR and normalized to WT levels in ΔE1 (A), ΔE2 (B), ΔE3 (C), ΔE1/2 (D) andΔE1/2/3 (E) cells under RA treatment for 24h. In (**A-E**), data are represented as min to max. The expression level in WT ESC was relatively set to 1 (As shown in blue dotted line). Indicated significance is based on Student’s *t*-test (Two-tailed, * indicates *P*< 0.05, ** indicates *P* < 0.01, *** indicates *P* < 0.001). In (**A-C, E**), n = 12, including 2 enhancer knockout cell lines, and 2 biological replicates per cell line with 3 technical replicates in 1 biological replicate. In (**D**), n = 6, including 1 enhancer knockout cell line, and 2 biological replicates per cell line with 3 technical replicates in 1 biological replicate. Source files are available in *Figure 6—source data 1*.

## DISCUSSION

The precise expressions of *HoxA* are extremely important for ESC differentiation. Although previous studies have shown that CBE plays an important role in the differentiation of ESC by regulating high-order chromatin structures (*Beagan et al., 2017; Narendra et al., 2015*), the identification of functional CBE and mechanisms of its regulation in *HoxA* expression remain largely unknown. Here, a new functional CBE^+47^ is identified in this study and a research model is proposed based on our results (*Figure 7*): In WT cells (Left), the three enhancers near CBE^+47^ acted as EEIC to interact with the 3’-end of *HoxA* chromatin to maintain *HoxA* normal expression, and then promote early correct differentiation of ESC induced by RA; in CBE^+47^-KO cells (Right), the absence of CBE^+47^ leads to reduced interactions between the three enhancers, thereby forming a relatively loose EEIC. Furthermore, the interactions between EEIC and central chromatin regions of *HoxA* are increased, thereby promoting over-expressions of *HoxA* and further enhancing RA-induced early ESC differentiation. These results indicate that a newly identified functional CBE^+47^ is required for maintaining the normal expressions of *HoxA* by organizing the precise interactions between its adjacent enhancers and *HoxA* chromatin, thereby regulating the early proper differentiation of ESC. Our findings highlight the unique and indispensable fine-regulatory roles of CBE as a functional element in high-order chromatin structures. The results also reveal direct regulatory effect of the CBE^+47^ on RA-induced *HoxA* expression and early ESC differentiation.

**Figure 7.**
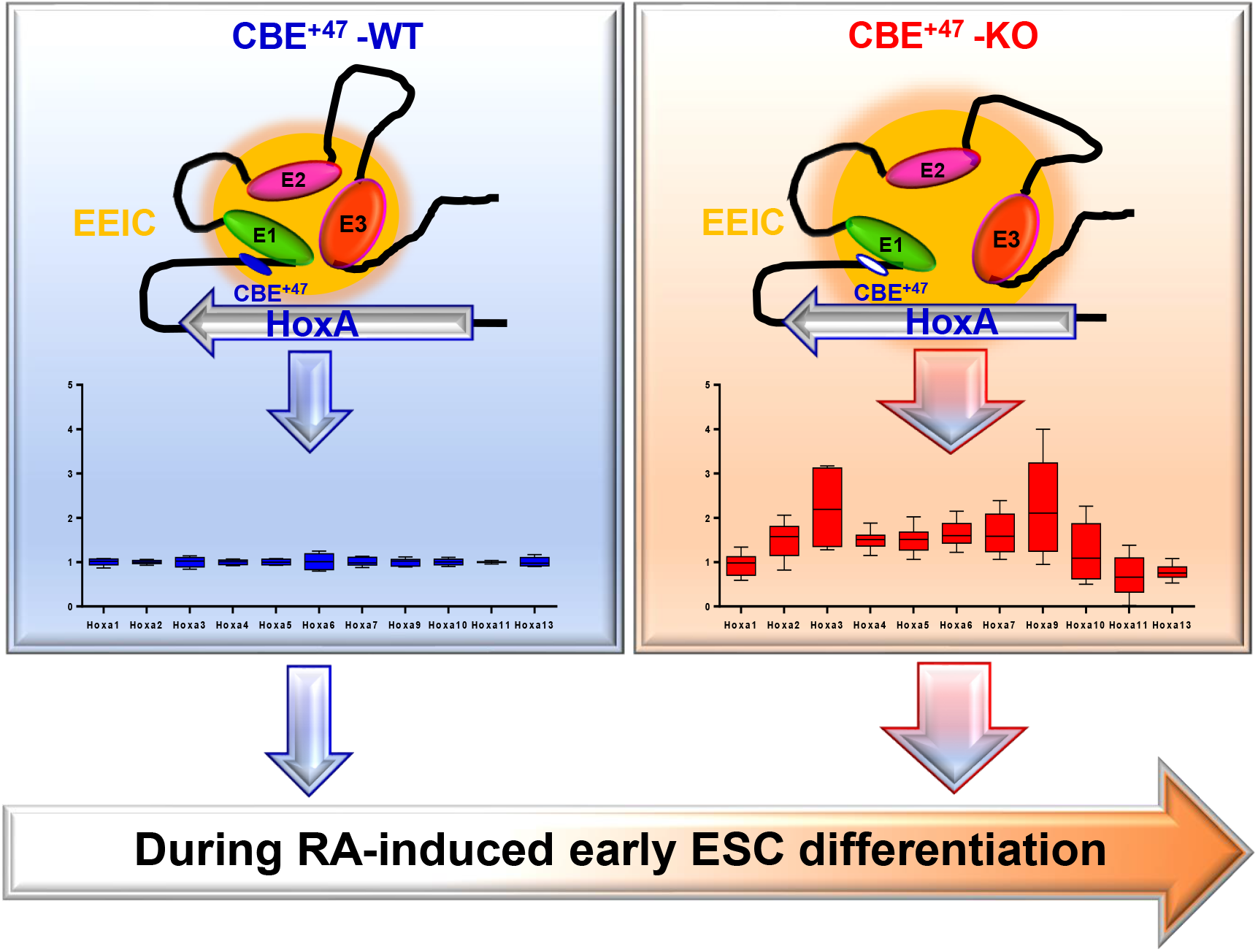
Proposed schematic illustration of the CBE^+47^ that regulates RA-induced *HoxA* expression and early ESC differentiation through orchestrating long-range chromatin interactions between its adjacent enhancers and *HoxA*. The CBE^+47^ working model is shown: in CBE^+47^-WT cells (Left), the three enhancers near CBE^+47^ acted as EEIC to interact with the 3’-end of *HoxA* chromatin to maintain *HoxA* normal expression, thereby promoting early proper differentiation of ESC induced by RA; in CBE^+47^-KO cells (Right), the absence of CBE^+47^ leads to reduced interactions between enhancers, thus forming a relatively loose EEIC, at the same time, interactions between EEIC and the central chromatin region of *HoxA* are increased, then promoting over-expressions of *HoxA*, further resulting in early ESC abnormal differentiation.

In an earlier study, Ferraiuolo MA, et al. have shown that *HoxA* has its special chromatin structures, which is significantly related to *HoxA* expression, and CTCF also plays a regulatory role in *HoxA* chromatin structures and gene expression (*Ferraiuolo et al., 2010*). Recent studies have focused on CBE within *HoxA* locus, including the TAD boundary, and found that the CBE could organize *HoxA* chromatin structures, maintain normal expressions of *HoxA*, and then regulate ESC differentiation (*Narendra et al., 2015*). The study of Varun Narendra, et al. showed that CBE5/6 knockout significantly promoted *HoxA* expression, changed *HoxA* chromatin structures and enhanced the neural differentiation of ESC (*Narendra et al., 2015*). Besides, studies in other cells, such as AML cells, have also revealed that CBE located at *HoxA* locus can significantly regulate expressions of *HoxA* and affect tumorigenesis (*Ghasemi et al., 2020; Huacheng Luo et al., 2018*). In this study, we find that besides several CBE in *HoxA* locus, there are other several CBE located downstream of *HoxA* within the same TAD (*Figure 1B*). Among them, the CBE^+47^ knockout significantly enhances RA-induced expressions of *HoxA* and promotes early ESC differentiation. These findings suggest that CBE^+47^ is a new functional CBE. However, several other CBE are still not studied, and whether these CBE have regulatory effects requires further research. In the recent studies, different research groups have developed a CRISPR-Cas9 based genetic screen method to detect functional CBE in breast cancer and AML (*Korkmaz et al., 2019; H. Luo et al., 2019*), which may be used to screen functional CBE in regulating ESC differentiation. These need further exploration.

In our subsequent transcriptome analysis, the results show that after CBE^+47^ deletion, genes in the same TAD with CBE^+47^ are significantly up-regulated compared with near TADs (*Figure 4–figure supplement 1*), which reveals that the CBE^+47^ may serve as a significant *cis*-regulatory element and play a local regulatory role. In addition, we also find that the RA signaling pathway-related genes are significantly up-regulated (Such as *Crabp2, Cyp26a1, Stra6*), while the pluripotent-related genes such as *Myc* and *Lin28a* are significantly down-regulated (*Figure 4–figure supplement 2*). These results suggest that the absence of CBE^+47^ significantly promotes RA-induced early ESC differentiation. Recent studies have shown that *Lin28a* can inhibit the expressions of *HoxA* and maintain limb development (*T. Sato et al., 2020*), while De Kumar B, et al. found that HOXA1 can bind to *Myc* near enhancers (*De Kumar et al., 2017*), which indicates that there may be positive feedback regulation between *HoxA* and *Myc, Lin28a*, etc. in early differentiation of ESC induced by RA. These speculations need further exploration, which will be helpful to understand the differentiation mechanisms and gene expression regulation in ESC.

RA can significantly induce *HoxA* expression during early ESC differentiation (*Figure 3–figure supplement 1*). Several studies have shown that at least three enhancers are needed for *HoxA* expression in early differentiation of ESC induced by RA (*Cao et al., 2017; Liu et al., 2016; Su et al., 2019; Yin et al., 2015*). The CBE^+47^ we studied is just between the three enhancers and *HoxA* (*Figure 5A*). As an insulator, CBE can isolate the interaction between enhancers and target genes and exert inhibitory effect (*Dowen et al., 2014; Gaszner & Felsenfeld, 2006; Kim et al., 2007*). Through our studies, first, we found that the three enhancers interact significantly as EEIC; second, the EEIC has significant interactions with *HoxA*; third, single enhancer interaction with *HoxA* also has its unique model. When the CBE^+47^ is deleted, we find that one is that the interactions between enhancers are significantly reduced, resulting in forming a relatively loose EEIC, which reveals that the CBE^+47^ is required for organizing these enhancers interaction; the second is that the interactions between enhancers and *HoxA* changes significantly, specifically, the interaction of E1 becomes more concentrated in the middle, and the interaction of E2 and E3 with *HoxA* also moves from the 3’-end to the middle regions. Furthermore, we also find that the deletion of the CBE^+47^ significantly promotes higher expressions of *Hoxa2-a9*, which is significantly corresponding to the changes of chromatin interactions between enhancers and *HoxA*. These findings reveal that CBE^+47^ can not only accurately organize interactions between enhancers, but also orchestrating interactions between enhancers and *HoxA*, thus ensuring the correct expressions of *HoxA*. Although the CBE^+47^ can organize high-order chromatin structures, and previous studies have shown that the binding direction of CTCF also can affect the formation of CTCF-mediated chromatin-loop (*de Wit et al., 2015*), other CBE paired with the CBE^+47^ to form chromatin-loop are unknown, which needs further exploration.

Previous studies have shown that multiple enhancers maintain RA-induced expressions of *HoxA* through long-range chromatin interactions with *HoxA* (*Cao et al., 2017; Liu et al., 2016; Su et al., 2019; Yin et al., 2015*). In this study, our results show that knockout of multiple enhancers can show significant synthesis regulation effect on *HoxA* expression (*Figure 6*), which is partly consistent with the previous report (*Cao et al., 2017*). Although these enhancers have been shown to have regulatory effects, we also find that several other enhancers located in the *Skap2* region in the same TAD with *HoxA* (*Figure 5A*). Whether these unreported enhancers have functional regulation effects needs to be further studied. Furthermore, our results also show that CBE^+47^ can significantly organize the interactions between enhancers (E1, E2 and E3). However, the datum shows that there are no significant CTCF-binding peaks in the E2 region, suggesting that other factors might be involved in regulating the interactions between enhancers. A recent study from Wang Ping, et al. have shown that RARα (Retinoic Acid Receptor alpha) can mediate chromatin interactions in AML cells by using the ChIA-PET method (Chromatin Interaction Analysis by Paired-End Tag sequencing) (*P. Wang et al., 2020*). Activation of target genes by RA signaling depends on RAR/RXR (retinoic acid receptor and retinoid X receptor) (*Rhinn & Dolle, 2012; Valcarcel, Holz, Jimenez, Barettino, & Stunnenberg, 1994*). Remarkably, we also find that RAR/RXR has significant binding peaks in E1, E2 and *HoxA* regions during RA-induced early ESC differentiation (data not shown) (*Simandi et al., 2016*), demonstrating that RAR/RXR may be involved in regulating these chromatin interactions, which needs further verification. Besides, we also find that chromatin interactions between the single enhancer and *HoxA* are different, and mechanisms of maintaining these different interactions need to be further studied, which will contribute to understanding the regulatory mechanisms of high-order chromatin structures on *HoxA* expression and ESC differentiation.

In summary, we identified a new functional CBE^+47^ that can regulate RA-induced expressions of *HoxA* and early ESC differentiation through orchestrating long-range chromatin interactions between its adjacent enhancers and *HoxA*. These findings can enhance our understanding of the intrinsic mechanisms of RA-induced early ESC differentiation, and further highlight the specific regulatory roles of CBE in high-order chromatin structures.

## MATERIALS AND METHODS

### Embryonic stem cells culture

Mouse ESC E14 was cultured as the previous scheme with a few modifications (*Su et al., 2019*). ESC were grown in culture dishes coated with 0.1% gelatin (Sigma, Lot # SLBQ9498V) in Dulbecco’s Modified Eagle’s Medium (DMEM, Gibco, Lot # 8119213) supplemented with 15% fetal bovine serum (FBS, AusGeneX, Lot # FBS00717-1), 1x nonessential amino acids (100x, Gibco, Lot # 2027433), 1x L-Glutamate (100x, Gibco, Lot # 2085472), 1x Penicillin-streptomycin (100x, Gibco, Lot # 2029632), 50 μM β-mercaptoethanol (Sigma, Cas # 60-24-2), 10 ng/mL LIF (ESGRO, Lot # ESG1107), 1 μM PD0325901 (a MEK inhibitor, MedChemExpress, Lot # HY-10254) and 3 μM CHIR99021 (a GSK inhibitor, MedChemExpress, Lot # HY-10182). The medium was replaced every 1-2 days. All cells are maintained at 37 °C in a 5% CO_2_ incubator.

### Retinoic acid (RA)-induced early ESC differentiation

To induce ESC differentiation, cells were gently washed with 1x phosphate buffer saline solution (PBS), dissociated and plated at an appropriate density on gelatin-coated plates in LIF/2i withdrawal medium supplemented with 2μm RA (Solarbio®, Lot # 1108F031). The culture medium was replaced at the given point-in-time.

### RNA extraction, reverse transcription, and quantitative Real-Time PCR (qRT-PCR)

Total RNA was extracted from differentiated or undifferentiated ESC using TRIzol Reagent (Life Technologies, Lot # 213504). cDNA synthesis was performed using a PrimerScriptTM RT reagent Kit with gDNA Eraser (TaKaRa, Lot # AJ51485A) according to the manufacturer’s instruction. PCR reactions were performed using HieffTMqPCR SYBR Green Master Mix (YEASEN, Lot # H28360) and a BioRad CFX Connect Real-Time system. PCR cycling conditions were as the previous protocol: 95 °C for 5 min, 40 cycles of 95 °C for 15 s, 60 °C for 15 s, and 72 °C for 30 s. And then, a melting curve of amplified DNA was subsequently acquired (*Su et al., 2019*). Quantification of target genes was normalized to *Gapdh* expression and experimental control through ΔΔCt methods (*Schmittgen & Livak, 2008*). Primer sequences used in this study are shown in *Supplementary file 1*.

### CRISPR/Cas9-mediated CBE^+47^ and enhancers deletion in ESC

The CRISPR/Cas9 system was used following published protocols (*Cong et al., 2013; Engreitz et al., 2016; Su et al., 2019*). Briefly, target-specific guide RNAs (sgRNAs) were designed using an online tool (http://chopchop.cbu.uib.no/). sgRNAs with the appropriate site and score were selected. sgRNA sequences are shown in *Supplementary file 2*. For CBE and enhancer knockout, sgRNAs were cloned into a Cas9-puro vector using the Bsmb1 site. ESC were transfected with two sgRNA plasmids using Lipofectamine 3000 (Life Technologies, Lot # 2125386), and 24 hours later, cells were treated with 5 μM puromycin (MCE, Lot # 64358) for 24 hours and then cultured in medium without puromycin for another 5 ~ 7 days. Individual colonies were picked and validated by genomic DNA PCR and subsequent Sanger DNA sequencing. Genotyping PCR primers are listed in *Supplementary file 3*.

### Alkaline Phosphatase (AP) Staining analysis in WT and CBE^+47^ deletion cells

Low-density ESC were planted in 12-well plates coated with gelatin for 4 days. Then, ESC were washed twice with PBS and incubated with AP Staining kit™ (Cat # AP100R-1, System Biosciences) following the manufacturer’s instructions. Digital images were taken by Olympus Inverted Fluorescence Microscope.

### Enhancer Capture-C and data analysis

The Capture-C probes were designed using an online tool within the three enhancers regions: (http://apps.molbiol.ox.ac.uk/CaptureC/cgi-bin/CapSequm.cgi) E1 bait (Mouse, mm10, chr6: 52012160 - 52015360), E2 bait (Mouse, mm10, chr6:52075358 - 52076639), E3 bait (Mouse, mm10, chr6: 52101884 - 52104217). Probe sequences are listed in *Supplementary file 4*.

Capture-C libraries were prepared as previously described with minor modifications (*Davies et al., 2016; Hughes et al., 2014; Su et al., 2019*). Briefly, RA-induced differentiated ESC (WT and CBE^+47^-KO) were fixed with 1% (vol/vol) formaldehyde for 10 min at room temperature, quenched with 125 mM glycine in PBS, and then lysed in cold lysis buffer [10 mM Tris-HCl, pH7.5, 10 mM NaCl, 5 mM MgCl_2_, 0.1 mM EGTA, 0.2% NP-40, 1× complete protease inhibitor cocktail (Roche, Lot # 3024150)]. Chromatins was digested with DpnII (New England Biolabs, Lot # 10014860) at 37°C overnight. Fragments were then diluted and ligated with T4 DNA ligase (Takara, Lot # 1211707) at 16°C overnight. Cross-linking was reversed by overnight incubation at 60°C with proteinase K (Bioline, Lot # BIO-37037). Then 3C libraries were purified by phenol-chloroform followed by chloroform extraction and ethanol-precipitated at −80°C overnight. Sequencing libraries were prepared from 10 μg of the 3C library by sonication to an average size of 200 ~ 300 bp and indexed using NEBnext reagents (New England Biolabs, Lot # 0031607), according to the manufacturer’s protocol. Enrichment of 2 μg of an indexed library incubated with 3 μM of a pool of biotinylated oligonucleotides was performed using the SeqCap EZ Hybridization reagent kit (Roche/NimbleGen, Lot # 05634261001), following the manufacturer’s instructions. Two rounds of capture employing 48~72 and 24h hybridizations, respectively, were used. Correct library size was confirmed by agarose gel electrophoresis, and DNA concentrations were determined using a Qubit 2.0 Fluorometer (Thermo Fisher Scientific, Lot # 0000248352). All sequencing was performed on Hi-Seq 2500 platforms using paired 150 bp protocols (Illumina).

Capture-C data were analyzed using the previously described methods (*Davies et al., 2016; Hughes et al., 2014; Su et al., 2019*). Briefly, the clean paired-end reads were reconstructed into single reads using FLASH (*Magoc & Salzberg, 2011*). After in silico DpnII digestion using the DpnII2E.pl script, the reads were mapped back to the mm10 mouse genome using Bowtie1. At last, chimeric reads containing captured reads and Capture-Reporter reads were analyzed using CCanalyser3.pl. The results can be visualized using the Integrated Genome Browser (IGV) (*Freese, Norris, & Loraine, 2016*) and online tool (https://epgg-test.wustl.edu/browser/) (*X. Zhou et al., 2013*).

### RNA-seq and bioinformatics analysis

ESC were lysed with Trizol reagent (Life Technologies, Lot # 265709) and RNA was extracted according to the manufacturer’s instructions. RNA was then sent to a sequencing company (Novogene, Tianjin, China) for sequencing. Clean reads were mapped to Ensemble mm10 mouse genome using Hisat2 with default parameters. Gene reads were counted by Htseq (*Anders, Pyl, & Huber, 2015*). Fold changes were computed as the log2 ratio of normalized reads per gene using the DEseq2 R package (*Love, Huber, & Anders, 2014*). Genes expression with | log2 (fold change) | ⩾ 1 (*P* < 0.05) were considered as significantly altered. Heatmaps were drawn with heatmap2. Two biological replicates were analyzed for each experimental condition.

### Gene Ontology (GO) and Kyoto Encyclopedia of Genes and Genomes (KEGG) pathway analyses

GO and KEGG pathway analyses were performed using the online tools: Metascape (http://metascape.org/gp/index.html#/main/step1) (*Y. Zhou et al., 2019*) and DAVID Functional Annotation Bioinformatics Microarray Analysis tool (http://david.abcc.ncifcrf.gov/) (*Huang, Sherman, & Lempicki, 2009*).

### Statistical analyses

Data were analyzed by Student’s *t*-test (Two-tailed) unless otherwise specified. Statistically significant *P*-values are indicated in Figures as follows: * indicates *P* < 0.05, ** indicates *P* <0.01, *** indicates *P* <0.001.

## Data availability

The data that support this study are available from the corresponding author upon request. The raw data reported in this manuscript for the DNA-seq and RNA-seq data have been deposited in NCBI Gene Expression Omnibus (GEO, https://www.ncbi.nlm.nih.gov/geo/) under the accession number **GSE154495**. The published datasets were download using online tools (http://cistrome.org/db/#/ and http://promoter.bx.psu.edu/hi-c/) and analyzed in this study (*Y. Wang et al., 2018; R. Zheng et al., 2019*).

## The following previously published data sets were used

NCBI Gene Expression Omnibus

1. Ali Shilatifard (2017)

ID GSM2588420. SET1A/COMPASS and shadow enhancers in the regulation of homeotic gene expression

NCBI Gene Expression Omnibus

2. Su G, Guo D, Lu W (2019)

ID GSE124306. A distal enhancer maintaining Hoxa1 expression orchestrates retinoic acid-induced early ESCs differentiation.

NCBI Gene Expression Omnibus

3. Bonev B, Mendelson Cohen N, Szabo Q, Fritsch L, Papadopoulos G, Lubling Y, Xu X, Lv X, Hugnot J, Tanay A, Cavalli G (2017)

ID GSE96107. Multi-scale 3D genome rewiring during mouse neural development.

NCBI Gene Expression Omnibus

4. Chieh-Chun Chen (2012)

ID GSM859491. Genome-wide analysis of K3K27ac, H2AZ and methylation data in mouse embryonic stem cells.

NCBI Gene Expression Omnibus

5. Ali Shilatifard (2018)

ID GSM2630487. An Mll4/COMPASS-Lsd1 epigenetic axis governs enhancer function and pluripotency transition in embryonic stem cells.

NCBI Gene Expression Omnibus

6. Pengfei Yu (2012)

ID GSM881353. Genome-wide analysis of histone modification, protein-DNA binding, cytosine methylation and transcriptome data in mouse and human ES cells and pig iPS cells.

NCBI Gene Expression Omnibus

7. Hendrik Marks (2014)

ID GSM1258237. Mll2 is required for H3K4 trimethylation on bivalent promoters in ES cells whereas Mll1 is redundant.

NCBI Gene Expression Omnibus

8. Lusy Handoko (2011)

ID GSM699165. CTCF-Mediated Functional Chromatin Interactome in Pluripotent Cells.

NCBI Gene Expression Omnibus

9. Richard A Young (2017)

ID GSM2645432. YY1 is a structural regulator of enhancer-promoter loops.

NCBI Gene Expression Omnibus

10. Richard A Young (2014)

ID GSM1439567. he developmental potential of iPSCs is greatly influenced by the selection of the reprogramming factors.

NCBI Gene Expression Omnibus

11. Richard A Young Consortium (2010)

ID GSM560345. Control of Embryonic Stem Cell State by Mediator and Cohesin.

NCBI Gene Expression Omnibus

12. Hendrik Marks (2014)

ID GSM1276711. Mll2 is required for H3K4 trimethylation on bivalent promoters in ES cells whereas Mll1 is redundant.

NCBI Gene Expression Omnibus

13. ENCODE DCC (2014)

ID GSM1014154. A comparative encyclopedia of DNA elements in the mouse genome.

NCBI Gene Expression Omnibus

14. Xiaoying Bai (2017)

ID GSM2651154. Transcription Pausing Regulates Mouse Embryonic Stem Cell Differentiation.

## ACKNOWLEDGEMENTS

We thank all members of our laboratory for many helpful discussions. This work was supported by the National Key R&D Program of China (NO.2017YFA0102600), Chinese National Natural Science Foundation (NSFC31530027).

## Author contributions

W.L., G.S. and L.Z. conceived and supervised the study and designed the experiments. G.S., J.C., M.L., J.Z., J.B., Z.Z. and J.S. performed the experiments and analyzed the data. J.C., D.G. and W.W. analyzed sequencing data. G.S. and W.W. performed bioinformatic analysis. G.S., W.L. and L.Z wrote the manuscript. All authors have read and approved the final manuscript.

---

**Figure 1—figure supplement 1. CBE^+47^ is located at the second intron of *Halr1* in ESC.**

(**A**) IGV (Integrative Genomics Viewer) screenshot showing refseq gene locus (*Halr1*).

(**B**) IGV screenshots showing gene tracks of CTCF, MED1, MED12 and YY1 ChIP-seq signal occupancy at *Halr1* locus in ESC.

(**C-D**) IGV view of selected ChIP-seq tracks at Halr1 locus in ESC. Shown are H3K27ac, H3K4me1, H3K4me2, H3K4me3, Pol ll, ATAC-seq, and DNase. Blue shadowing indicates the CBE^+47^ region.

**Figure 3—figure supplement 1. qRT-PCR analyses *HoxA* expression during RA-induced ESC differentiation.**

qRT-PCR results showing expressions of *HoxA* in ESC after RA treatment (0h, 12h, 24h and 48h). The expression level in untreated ESC (0h) was relatively set to 1. Data are represented as mean ± SD. N = 6, including 1 cell line (ESC), and 2 biological replicates per cell line with 3 technical replicates in 1 biological replicate.

Source files are available in *Figure 3—figure supplement 1—source data 1*.

**Figure 3—figure supplement 2. qRT-PCR analyses expressions of *HoxA*-related lncRNAs in WT and CBE^+47^-KO cells during RA-induced differentiation.**

(**A**) Diagram showing RA-induced WT and CBE^+47^-KO cells differentiation.

(**B-D**) qRT-PCR analyses *HoxA*-related lncRNAs expression in WT and CBE^+47^-KO cells during RA-induced differentiation.

In (**B-D**), data are represented as min to max. The expression level in WT ESC was relatively set to 1 (As shown in blue dotted line). Indicated significance is based on Student’s *t*-test (Two-tailed, * indicates *P* < 0.05, ** indicates *P* < 0.01, *** indicates *P* < 0.001).

In (**B-D**), n = 12, including 2 CBE^+47^-KO cell lines (CBE^+47^-KO1# and CBE^+47^-KO2#), and 2 biological replicates per cell line with 3 technical replicates in 1 biological replicate.

Source files are available in *Figure 3—figure supplement 2—source data 1*.

**Figure 4—figure supplement 1. CBE^+47^ served as a *cis*-regulatory element regulates gene expression within the same TAD.**

(**A**) Hi-C interaction map of the ~ 2.0 Mb region surrounding the *HoxA* in undifferentiated ESC. Data were extracted from Bonev B et al. 2017.

(**B**) RNA-seq results shown the Heatmap of *HoxA*.

(**C-D**) data show that CBE^+47^ as cis-regulatory element regulates gene expression within the same TAD.

In (**D**), NS: not significant. Indicated significance is based on Student’s t-test (Two-tailed, * indicates *P* < 0.05, ** indicates *P* < 0.01, *** indicates *P* < 0.001).

Source files are available in *Figure 4—figure supplement 1—source data 1*.

**Figure 4—figure supplement 2.** RNA-seq results shown the Heatmap of *HoxB/C/D* cluster genes (**A**), RA signaling-(**B**) and pluripotency- (**C**) related genes.

**Figure 4—figure supplement 3. Down-regulated and up-regulated genes to enriched GO-BP clusters.**

**Figure 6—figure supplement 1. CRISPR-cas9 mediated enhancers knockout.**

(**A**) Schematic showing CRISPR/Cas9-mediated deletion of the enhancers using specially paired sgRNAs (Red silk arrows). Indicated primers (Blue silk arrows) were used to distinguish enhancers knockout from wild type clones.

(**B**) Validation of knockout cell lines by genomic DNA PCR. Agarose gel electrophoresis images from representative clones. The red dashed boxes mark the target genomic DNA PCR products. ΔE3 cell lines have been reported in our previous study (Su et al., 2019).

**Figure 6—figure supplement 2. CBE^+47^ and enhancers regulate the *HoxA* expression.**

(**A**) The data show that CBE^+47^ and enhancers have inhibitory and enhanced regulatory effects on *HoxA* expression, respectively.

(**B**) Comparing the difference in regulation of *HoxA* between enhancers. Indicated significance (*P*-value) is based on Student’s *t*-test (Two-tailed).

Source files are available in *Figure 6—figure supplement 2—source data 1*.

## FIGURE SUPPLEMENTS

**Figure 1—figure supplement 1. CBE^+47^ is located at the second intron of Halr1 in ESC.**

**Figure 4—figure supplement 2.** RNA-seq results shown the Heatmap of HoxB/C/D cluster genes (**A**), RA signaling-(**B**) and pluripotency- (**C**) related genes.

**Figure 4—figure supplement 3. Down-regulated and up-regulated genes to enriched GO-BP clusters.**

**Figure 6—figure supplement 1. CRISPR-cas9 mediated enhancers knockout.**

**Figure 6—figure supplement 2. CBE^+47^ and enhancers regulate the *HoxA* expression.**

## SUPPLEMENTARY FILES

**Supplementary file 1. Sequences of qRT-PCR primers (5’-3’) used in this study**

**Supplementary file 2. sgRNAs used in this study**

**Supplementary file 3. Primers were used to identify the deletion clones in this study**

**Supplementary file 4. Synthetic oligos for Capture-C (5’-3’) (5’-biotin)**

## SOURCE DATA FILES

**Figure 2—source data 1.xlsx**

**Figure 3—source data 1.xlsx**

**Figure 4—source data 1.xlsx**

**Figure 6—source data 1.xlsx**

**Figure 3—figure supplement 1—source data 1.xlsx**

**Figure 3—figure supplement 2—source data 1.xlsx**

**Figure 4—figure supplement 1—source data 1.xlsx**

**Figure 6—figure supplement 2—source data 1.xlsx**

